# Crystal structure of GH43 exo-β-1,3-galactanase from the basidiomycete *Phanerochaete chrysosporium* provides insights into the mechanism of bypassing side chains

**DOI:** 10.1101/2020.09.23.310037

**Authors:** Kaori Matsuyama, Naomi Kishine, Zui Fujimoto, Naoki Sunagawa, Toshihisa Kotake, Yoichi Tsumuraya, Masahiro Samejima, Kiyohiko Igarashi, Satoshi Kaneko

## Abstract

Arabinogalactan proteins (AGPs) are functional plant proteoglycans, but their functions are largely unexplored, mainly because of the complexity of the sugar moieties, which are generally analyzed with the aid of glycoside hydrolases. In this study, we solved the apo and liganded structures of exo-β-1,3-galactanase from the basidiomycete *Phanerochaete chrysosporium* (*Pc*1,3Gal43A), which specifically cleaves AGPs. It is composed of a glycoside hydrolase family 43 subfamily 24 (GH43_sub24) catalytic domain together with a carbohydrate-binding module family (CBM) 35 binding domain. GH43_sub24 lacks the catalytic base Asp that is conserved among other GH43 subfamilies. Crystal structure and kinetic analyses indicated that the tautomerized imidic acid function of Gln263 serves instead as the catalytic base residue. *Pc*1,3Gal43A has three subsites that continue from the bottom of the catalytic pocket to the solvent. Subsite -1 contains a space that can accommodate the C-6 methylol of Gal, enabling the enzyme to bypass the β-1,6-linked galactan side chains of AGPs. Furthermore, the galactan-binding domain in CBM35 has a different ligand interaction mechanism from other sugar-binding CBM35s. Some of the residues involved in ligand recognition differ from those of galactomannan-binding CBM35, including substitution of Trp for Gly, which affects pyranose stacking, and substitution of Asn for Asp in the lower part of the binding pocket. *Pc*1,3Gal43A WT and its mutants at residues involved in substrate recognition are expected to be useful tools for structural analysis of AGPs. Our findings should also be helpful in engineering designer enzymes for efficient utilization of various types of biomass.

## Introduction

Arabinogalactan proteins (AGPs) are proteoglycans characteristically localized in the plasma membrane, cell wall, and intercellular layer of higher land plants (1), in which they play functional roles in growth and development (2). The carbohydrate moiety of AGPs is composed of a β-1,3-D-galactan main chain and β-1,6-D-galactan side chain, decorated with arabinose, fucose, and glucuronic acid residues (1, 2). The chain lengths and frequencies of side chains are different among plant species, organs, and stages of development (3), and the overall structures of the carbohydrate moieties of AGPs are not yet fully understood. Degradation of polysaccharides using specific enzymes is one approach to investigate their structures and roles. In this context, exo-β-1,3-galactanase (EC 3. 2. 1. 145) specifically cleaves the non-reducing end β-1,3-linked galactosyl linkage of β-1,3-galactans to release D-galactose (Gal). In particular, it releases β-1,6-galactooligosaccharides together with Gal from AGPs (4, 5), and is therefore useful for structural analysis of AGPs.

The basidiomycete *Phanerochaete chrysosporium* produces an exo-β-1,3-galactanase (*Pc*1,3Gal43A; GeneBank accession No. BAD98241) that degrades the carbohydrates of AGPs when grown with β-1,3-galactan as a carbon source (6). *Pc*1,3Gal43A consists of a glycoside hydrolase (GH) family 43 subfamily 24 (GH43_sub24) catalytic domain and a carbohydrate-binding module (CBM) belonging to family 35 (designated as *Pc*CBM6 in the previous paper) based on the amino acid sequences in the Carbohydrate-Active enZymes (CAZy) database (http://www.cazy.org; 6-8). The properties of the enzyme have been analyzed using recombinant *Pc*1,3Gal43A expressed in the methylotrophic yeast *Pichia pastoris* (6). The CBM35 of *Pc*1,3Gal43A was characterized as the first β-1,3-galactan binding module, and *Pc*1,3Gal43A showed typical GH43_sub24 activity. The enzyme cleaves only β-1,3-linkages of oligosaccharides and polysaccharides, but produces β-1,6-galactooligosaccharides together with Gal. Thus, *Pc*1,3Gal43A specifically recognizes β-1,3-linked Gal, but can accommodate β-1,6-bound side chains (6).

Glycoside hydrolases are classified into families based on sequence similarity, while they are also divided into major two groups according to their catalytic mechanisms, i.e., inverting enzymes and retaining enzymes (9, 10). Inverting enzymes typically utilize two acidic residues that act as an acid and a base, respectively, and a hydroxyl group connected to anomeric carbon inverts from the glycosidic linkage after the reaction. GH43 enzymes are members of the inverting group, and share conserved Glu and Asp as the catalytic acid and base, respectively (8), but GH43_sub24 enzymes lack the catalytic base Asp (8, 11, 12). In *Ct*1,3Gal43A (from *Clostridium thermocellum*), Glu112 was thought to be the catalytic base (13), but in BT3683 (from *Bacteroides thetaiotamicron*), Glu367 (corresponding to Glu112 of *Ct*1,3Gal43A) was found not to act as a base, but to be involved in recognition of the C-4 hydroxyl group of the non-reducing terminal Gal, and instead, Gln577 is predicted to be the catalytic base in the form of an unusual tautomerized imidic acid (12). An example of GH lacking a catalytic base, endoglucanase V from *P. chrysosporium* (*Pc*Cel45A), is already known, and based on the mechanism proposed for this enzyme, it is possible that tautomerized Gln functions as a base in GH43_sub24, or that this Gln stabilizes nucleophilic water. *Pc*Cel45A lacks the catalytic base Asp that is conserved in other GH45 subfamilies (14), but it uses the tautomerized imidic acid of Asn as the base, as indicated by neutron crystallography (15). However, it is difficult to understand the situation in GH43_sub24, since no holo structure with a ligand at the catalytic center has yet been solved in this family. Moreover, no structure of eukaryotic GH43_sub24 has yet been reported.

The CBM35 module is composed of approximately 140 amino acids. This family includes modules with various binding characteristics, and decorated with xylans, mannans, β-1,3-galactans, and glucans (16–21). The family members are divided into four clusters based on their sequences and binding specificities (17). The structures of CBM35s binding with xylan, mannan, and glucan have already been solved (16–21), but no structure of β-1,3-galactan-binding CBM35 has yet been reported.

In the present manuscript, we solved the apo and liganded structures of *Pc*1,3Gal43A. Based on the results, we discuss the catalytic mechanism and the mode of ligand binding to CBM35 in the two-domain structure.

## Results

### Overall structure of *Pc*1,3Gal43A

The crystal structure of the selenomethionine (SeMet) derivative of *Pc*1,3Gal43A was first determined by means of the multiwavelength anomalous dispersion method, and this was followed by structure determination of the ligand-free wild-type (WT), the WT bound with Gal (WT_Gal), the E208Q mutant co-crystallized with β-1,3-galactotriose (Gal3; E208Q_Gal3), and the E208A mutant co-crystallized with Gal3 (E208A_Gal3). Data collection statistics and structural refinement statistics are summarized in Tables 1 and 2, respectively.

**Table 1.**
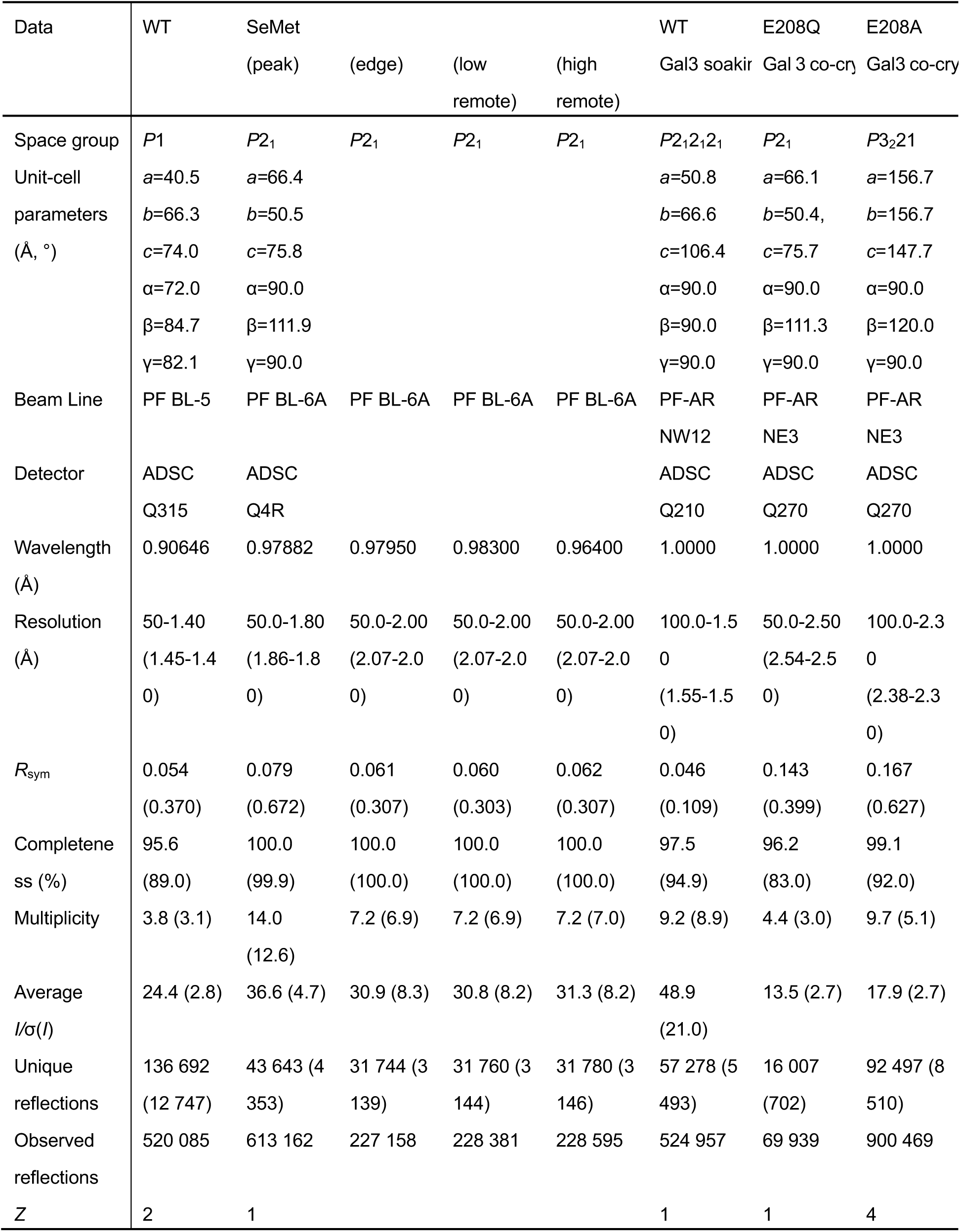
Data collection statistics

**Table 2.** Refinement statistics

| Data | WT | WT_Gal | E208Q_Gal3 | E208A_Gal3 |
| --- | --- | --- | --- | --- |
| Resolution range | 7.997 - 1.398<br>(1.448 - 1.398) | 41.56 - 1.500<br>(1.554 - 1.500) | 29.79 - 2.499<br>(2.588 - 2.499) | 30.66 - 2.300<br>(2.382 - 2.300) |
| Completeness (%) | 95.46 (87.82) | 97.51 (94.80) | 96.41 (85.67) | 98.78 (92.17) |
| Wilson B-factor | 12.76 | 10.11 | 29.91 | 30.40 |
| Reflections used in refinement | 136655 (12497) | 57105 (5474) | 15762 (1381) | 92011 (8507) |
| Reflections used for R-free | 6862 (630) | 2884 (272) | 799 (64) | 4568 (441) |
| R-work (%) | 15.47 (22.50) | 13.43 (12.71) | 16.62 (25.54) | 16.10 (22.39) |
| R-free (%) | 18.56 (26.28) | 16.00 (17.93) | 24.39 (42.53) | 21.43 (28.28) |
| Number of non-hydrogen atoms | 7966 | 3923 | 3576 | 14570 |
| macromolecules | 6615 | 3290 | 3235 | 12886 |
| ligands | 109 | 121 | 114 | 678 |
| solvent | 1242 | 512 | 227 | 1006 |
| Protein residues | 2106 | 427 | 428 | 1708 |
| RMS (bonds) | 0.008 | 0.006 | 0.008 | 0.011 |
| RMS (angles) | 1.22 | 0.87 | 0.94 | 1.05 |
| Ramachandran favored (%) | 97.29 | 97.41 | 94.13 | 95.76 |
| Ramachandran allowed (%) | 2.71 | 2.59 | 5.87 | 4.24 |
| Ramachandran outliers (%) | 0 | 0 | 0 | 0 |
| Rotamer outliers (%) | 0.81 | 0.55 | 0.29 | 0.36 |
| Clash score | 2.06 | 1.95 | 6.94 | 3.50 |
| Average B-factor ($\text{\AA}^2$) | 17.21 | 12.45 | 30.48 | 32.98 |
| macromolecules | 14.97 | 10.57 | 29.77 | 31.60 |
| ligands | 29.38 | 23.33 | 52.26 | 56.11 |
| solvent | 28.09 | 22.02 | 29.74 | 35.03 |
| PDB ID | 7BYS | 7BYT | 7BYV | 7BYX |

The recombinant *Pc*1,3Gal43A molecule is composed of a single polypeptide chain of 428 amino acids (Gln21-Tyr448) with two extra amino acids, Glu19 and Phe20, derived from the restriction enzyme cleavage site, which are disordered and thus were not observed. The protein is decorated with *N*-glycans, since it was expressed in *Pichia* yeast. Up to three sugar chains are attached at Asn79, Asn194, and Asn389; the attached chains vary in position and structure, and most contain one or two *N*-acetylglucosamine moieties.

*Pc*1,3Gal43A is composed of two domains, and ligands introduced by soaking or co-crystallization are located in a subsite of the catalytic domain or the binding site of CBM35 (Fig. 1). The N-terminal catalytic domain consists of a five-bladed b-propeller (Gln21-Gly325), as in other GH clan-F enzymes, and the C-terminal domain (*Pc*CBM35) takes a β-jellyroll fold (Thr326-Tyr448) structure, as in previously reported CBM35s (16–25). *Pc*CBM35 contains one calcium ion near the end of the first b-strand on a different domain surface from the plane to which the ligand binds (Fig. 1). The structure of *Pc*CBM35 is similar to those of other known CBM35s. The interface area is 686 A^2^ and includes many water molecules. The PDBePISA server (http://www.ebi.ac.uk/msd-srv/prot_int/cgi-bin/piserver) indicates that the enzyme forms a complex in the crystal, but this is an effect of crystallization, and the enzyme exists as a monomer in solution (data not shown).

**Fig. 1.**
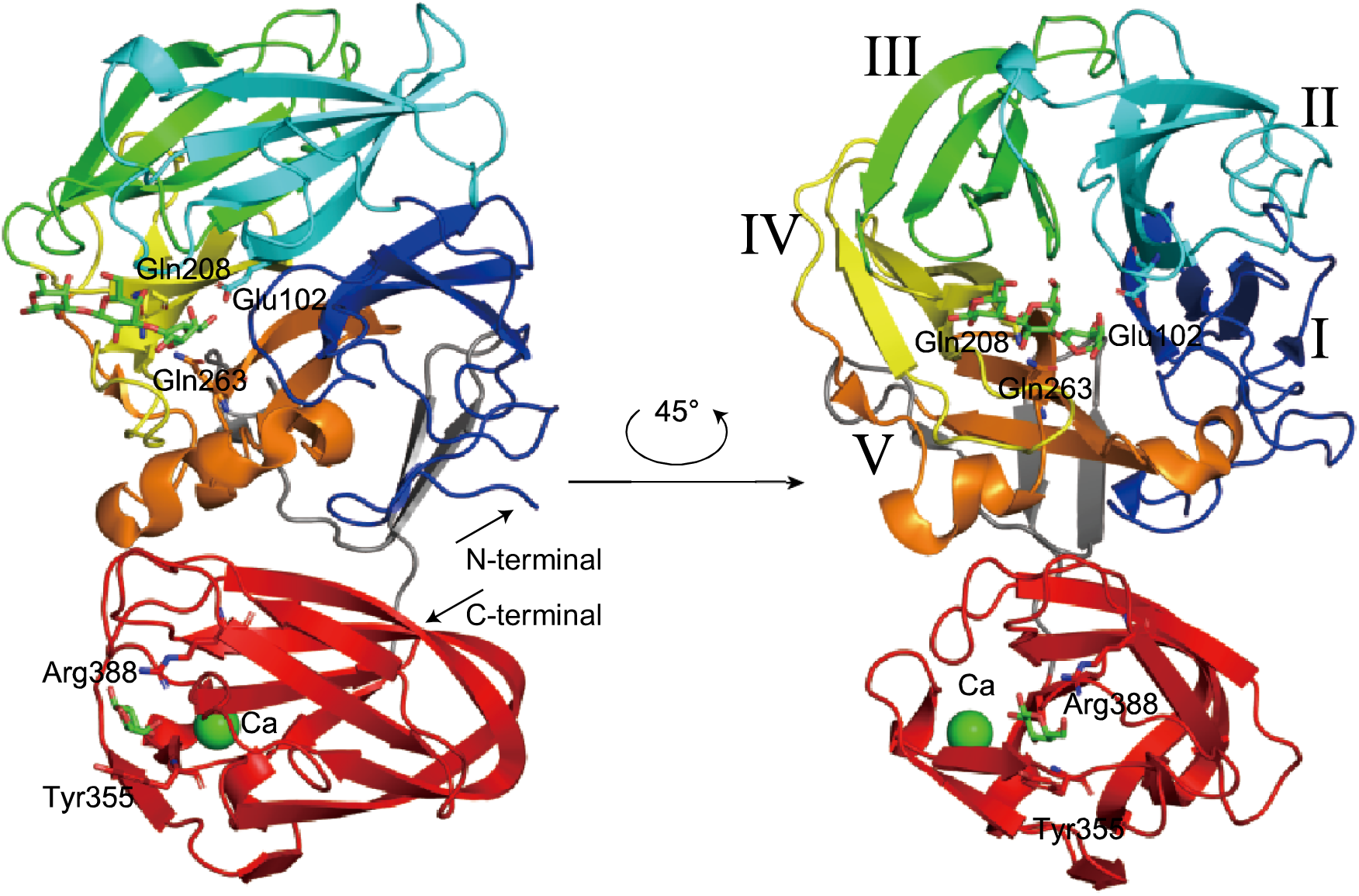
Overall structure of *Pc*1,3Gal43A. In the three-dimensional structure of *Pc*1,3Gal43A, the five blades of the catalytic domain are shown in blue (Gln21-Leu87), cyan (Ser88-Asp155), green (Ser156-Gly204), yellow (Ala205-Ser247), and orange (Ala248-Asp297) with successive roman numerals. The CBM (The326-Val448) is shown in orange. The linker connecting the two domains (Phe298-Gly325) is shown in gray.

### Sugar-binding structure of the *Pc*1,3Gal43A catalytic domain

The five-bladed b-propeller exhibits an almost spherical structure, and two central cavities are located at the ends of the pseudo-5-fold axis (Fig. 1). One of them contains the catalytic site and it is common in almost all GH43 enzymes. The catalytic site is located in the center of the five-bladed b-propeller, whose blades are formed by Gln21 or Asn22-Leu87 (I in Fig. 1), Ser88-Asp155 (II in Fig. 1), Ser156-Gly204 (III in Fig. 1), Ala205-Ser247 (IV in Fig. 1), and Ala248-Asp297 (V in Fig. 1).

As shown in Fig. 2, the Gal3 molecule co-crystallized with the E208Q mutant occupies subsites -1, +1, and +2 of the catalytic site, from the non-reducing end to the reducing end. Gal_-1_ is located at the bottom of the catalytic cavity, and Gal_+1_ and Gal_+2_ extend linearly outwards. Gal_+1_ is half buried in the cavity, whereas Gal_+2_ is exposed at the surface (Fig. 2A).

**Fig. 2.**
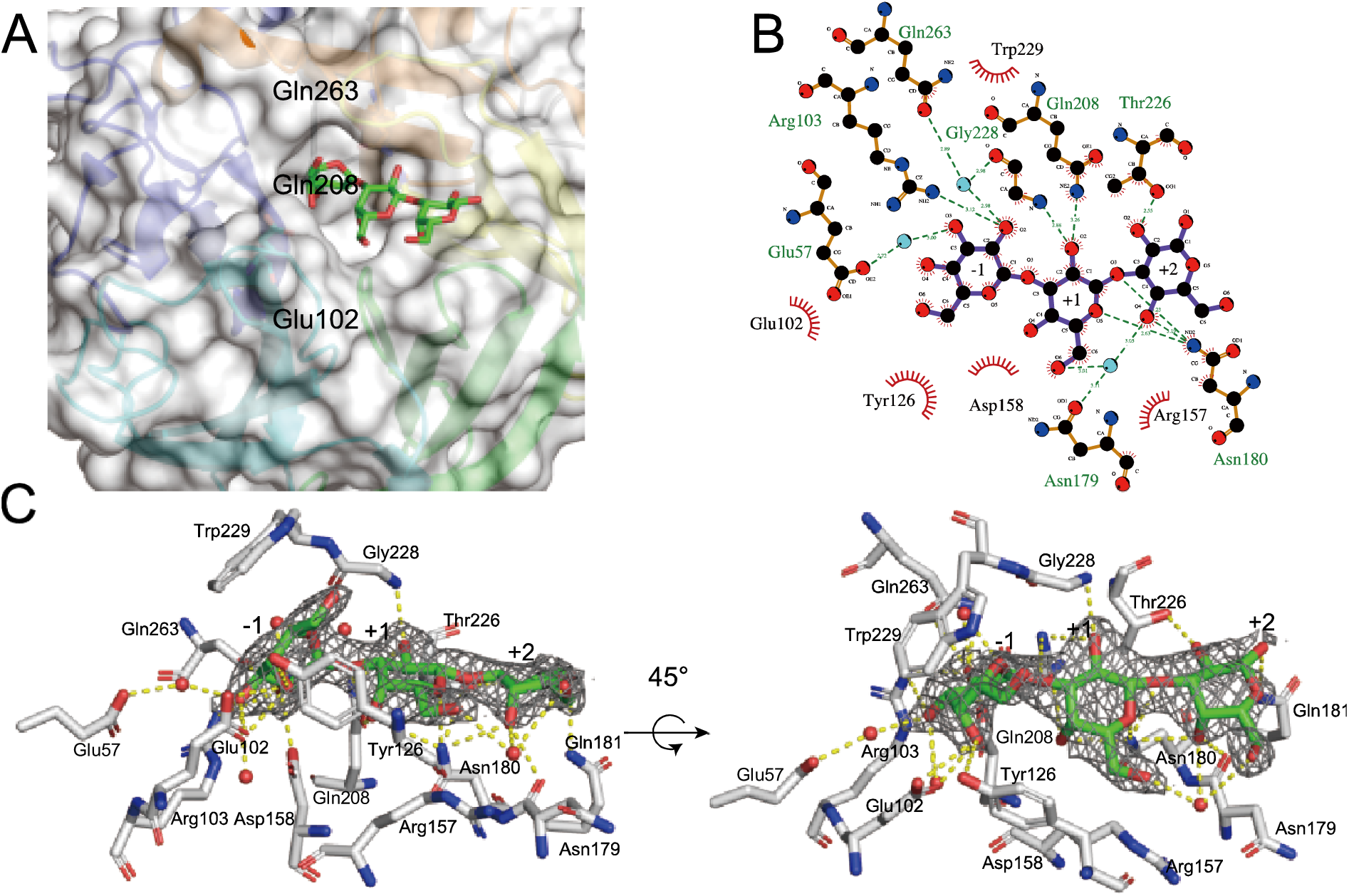
Gal3 binding mode at the catalytic site. A: The surface structure of the catalytic center. Gal3 is represented as green (carbon) and red (oxygen) sticks. B: Schematic diagram showing the interaction mode at the catalytic center. Black, red, and blue show carbon, oxygen, and nitrogen, respectively. Red lines indicate the hydrophobically interacting residues. This diagram was drawn with LigPlot+ (version 1.4.5). C: The 2Fo-Fc omit map is drawn as a blue mesh (0.8 sigma). Residues are shown in white (carbon), red (oxygen), and blue (nitrogen). Gal3 is shown in green (carbon) and red (oxygen). Yellow dots indicate hydrogen bonds and/or hydrophobic interaction and red spheres show water molecules interacting with ligands or residues.

Gal_-1_ adopts a *^1^S_3_* skew boat conformation and interacts with many residues via hydrogen bonds and hydrophobic interactions. As shown in Fig. 2B and C, the C-2 hydroxyl group of Gal_-1_ forms hydrogen bonds with NH_2_ of Arg103 and with OE1 of Gln263 via water. In addition, this water molecule is bound with O of Gly228. The C-3 hydroxyl group of Gal_-1_ also forms a hydrogen bond with OE2 of Glu57 via water. Glu102, Tyr126, Asp158, Gln208, Thr226, Trp229, and Gln263 interact with Gal3 through hydrophobic interactions. Notably, Trp229 supports the flat C3-C4-C5-C6 structure of Gal_-1_, and Tyr126 recognizes the C-6 methylol and C-4 hydroxyl groups, while Glu102 recognizes the C-3 hydroxyl and C-4 hydroxyl groups. In Gal_+1_ (as shown in Fig. 2B and C), the C-2 hydroxyl group forms a hydrogen bond with NE2 of Gln208 and N of Gly228, while O5 forms a hydrogen bond with ND2 of Asn180, and C-6 hydroxyl group forms a hydrogen bond with OD1 of Asn179 via water. Tyr126, Arg157, Asn180, and Gln208 interact hydrophobically with Gal. In Gal_+2_ (Fig. 2B and C), the C-2 and C-4 hydroxyl groups form hydrogen bonds with OG1 of Thr226 and ND2 of Asn180, respectively. In addition, Thr226 interacts with Gal_+2_ through hydrophobic interaction. Furthermore, the glycosidic oxygen between Gal_+1_ and Gal_+2_ interacts with ND2 of Asn180 through a hydrogen bond.

In the structure of WT_Gal, one Gal was found at subsite -1, taking a ^4^*C*_1_ chair conformation with α-anomeric conformation of the C-1 hydroxyl group (data not shown). The binding mode of Gal_-1_ is almost the same as that in E208Q_Gal3, but the C-1 hydroxyl group in the axial position forms hydrogen bonds with Gly228 and Gln263. No Gal3 molecule was observed at the catalytic domain in the structure of the Gal3 co-crystallized E208A mutant.

In order to identify the catalytic residues, we examined the relative activity of WT and the six mutants towards β-1,3-galactobiose (Gal2) and Gal3. WT showed 5.58±0.35 and 11.15±0.39 units of activity (μmol Gal /min/nmol enzyme) towards Gal2 and Gal3, respectively, whereas the six mutants showed no detectable activity (Fig. S1), suggesting that these residues are all essential for the catalysis.

### Sugar-binding structure of CBM35 in *Pc*1,3Gal43A

*Pc*1,3Gal43A has one CBM35 domain at the C-terminus. We previously reported that this enzyme has a CBM6-like domain (6), but it has been reclassified into the CBM35 family (7). The β-jellyroll fold domain is accompanied by a single calcium ion binding site on a different domain surface than the surface to which the ligand at the end of the first β chain binds, and this corresponds to a conserved calcium ion binding site in CBM35s. Some CBM35 modules bind another calcium ion at a site at the top of domain (16), but *Pc*CBM35 lacks this second calcium ion binding site (Fig. 1).

In E208A_Gal3, electron density of Gal3 was observed in the ligand binding site of *Pc*CBM35. As illustrated in Fig. 3A and S2, 2Fo-Fc omit maps showed that the binding mode of *Pc*CBM35 with ligands is “exo-type”, corresponding to type-C CBM (26). The asymmetric unit of E208A_Gal3 contained four *Pc*1,3Gal43A molecules and each molecule binds to the non-reducing end of Gal3 (called Gal_site 1), as in other CBM35 modules. However, the middle Gal (Gal_site 2) and the reducing end Gal (Gal_site 3) are found in two main locations (Fig. 3), though residues involved in the interactions with the ligand in each molecule were mostly shared. The Gal_site 1 forms hydrogen bonds with Tyr355 and Arg388 and interacts hydrophobically with Leu342, Gly354, Tyr438, and Asp441. The Gal_site 2 interacts hydrophobically with Gly383 and Asp384. The main ligand interaction in the Gal_site 3 involves Gly409 and Gly410, but in addition to these residues, Asn411 is also involved in ligand recognition in chain C (Fig. 4).

**Fig. 3.**
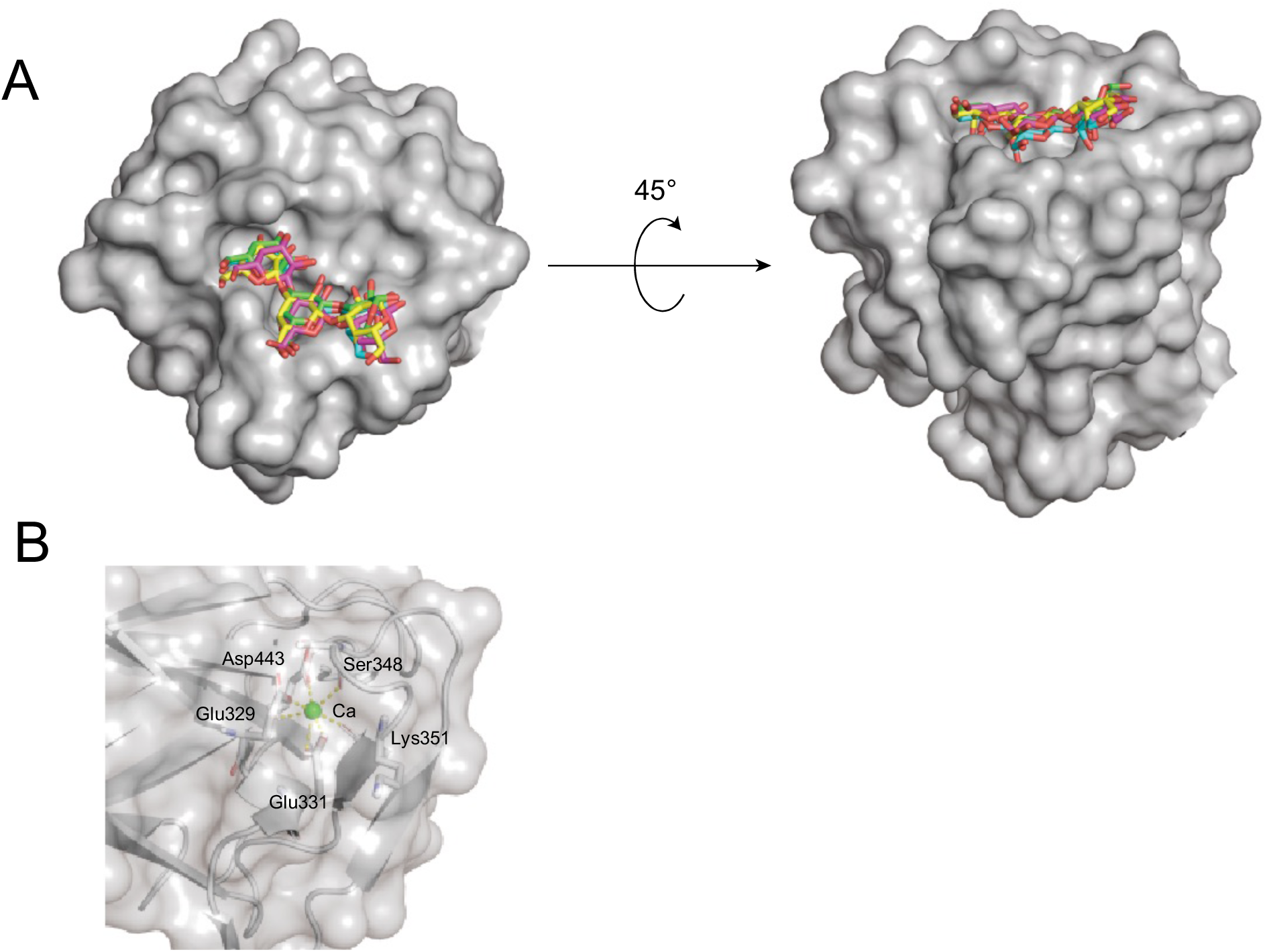
Surface structures of the CBM. A: Substrate binding mode at CBM35. Green, cyan, magenta, and yellow indicate carbons of chains A, B, C, and D of E208A_Gal3, respectively, and red shows oxygen. The left side is the non-reducing end of Gal3, and the right side is the reducing end. B: calcium ion binding mode at CBM35. Calcium ion is represented as green spheres and interacting residues are shown as stick models. Yellow dots indicate interaction.

**Fig. 4.**
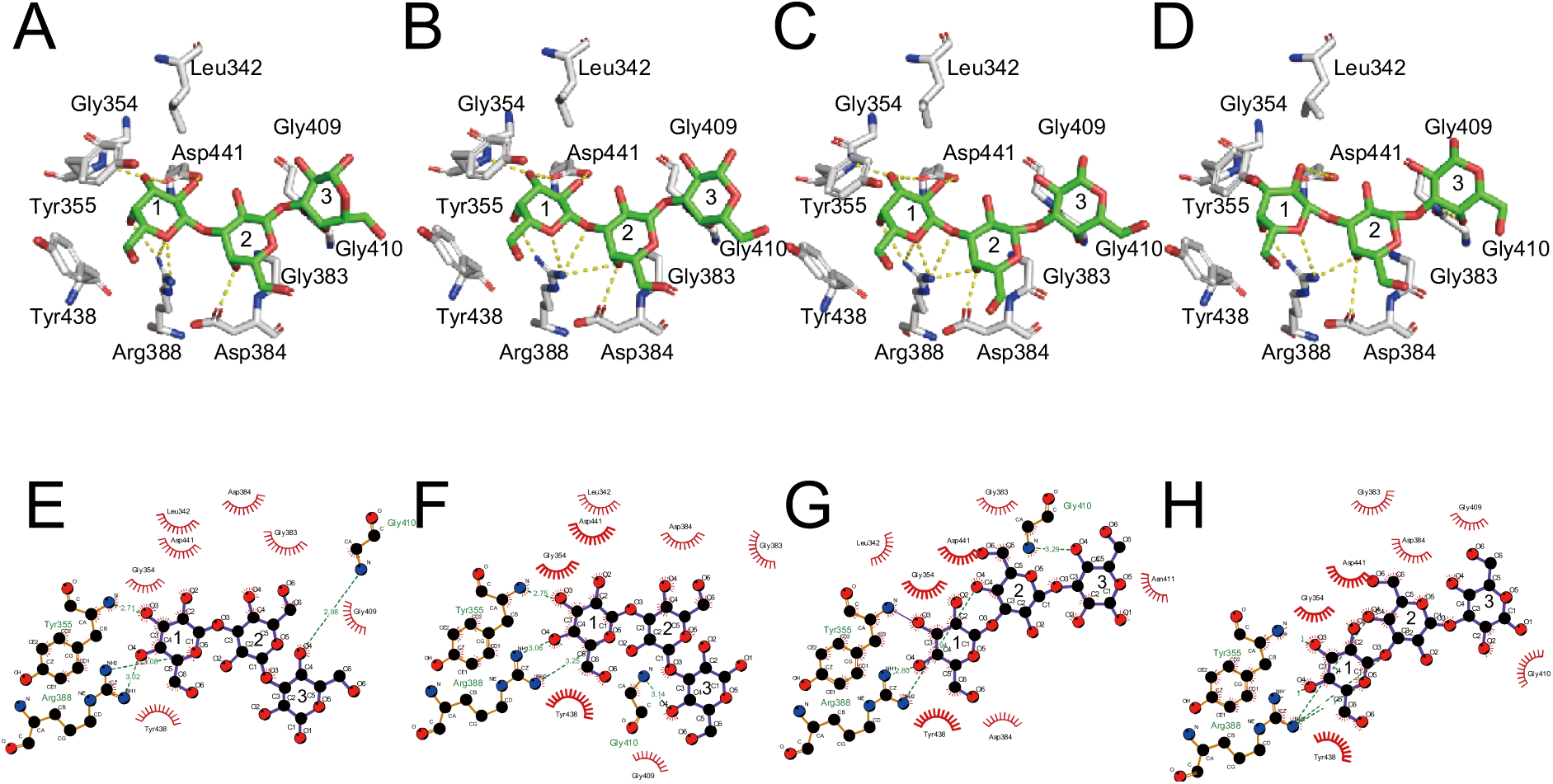
Ligand interaction mode at CBM35. A and E, B and F, C and G, and D and H are chains A, B, C, and D of E208A, respectively. A to D: Interaction modes between ligand and CBM35 residues. Atoms are indicated in the same colors as in Fig. 2. E to G: Schematic diagram showing the interaction mode at CBM35. Atoms are indicated in the same colors as in Fig. 2. Sugar binding sites are named Gal_site 1, Gal_site 2, and Gal_site 3 from the non-reducing end of the sugar, and in this figure, they are labeled 1, 2, and 3, respectively.

### Ensemble refinement

In order to understand the fluctuation of ligands, ensemble refinements were performed with the refined models. This method produces ensemble models by employing the combination of X-ray structure refinement and molecular dynamics. These models can simultaneously account for anisotropic and anharmonic distributions (27). Four different pTLS values (%) of 0.6, 0.8, 0.9, and 1.0 were set for each model. Table 3 shows the statistical scores of the refinement with the most appropriate pTLS value for each model. Focused views of the catalytic site in the catalytic domain and the ligand binding site of the CBM are shown in Fig. 5 and 6, respectively. Note that structures containing multiple molecules in the asymmetric unit (WT and E208A_Gal3) are found for all molecules in this paper.

**Fig. 5.**
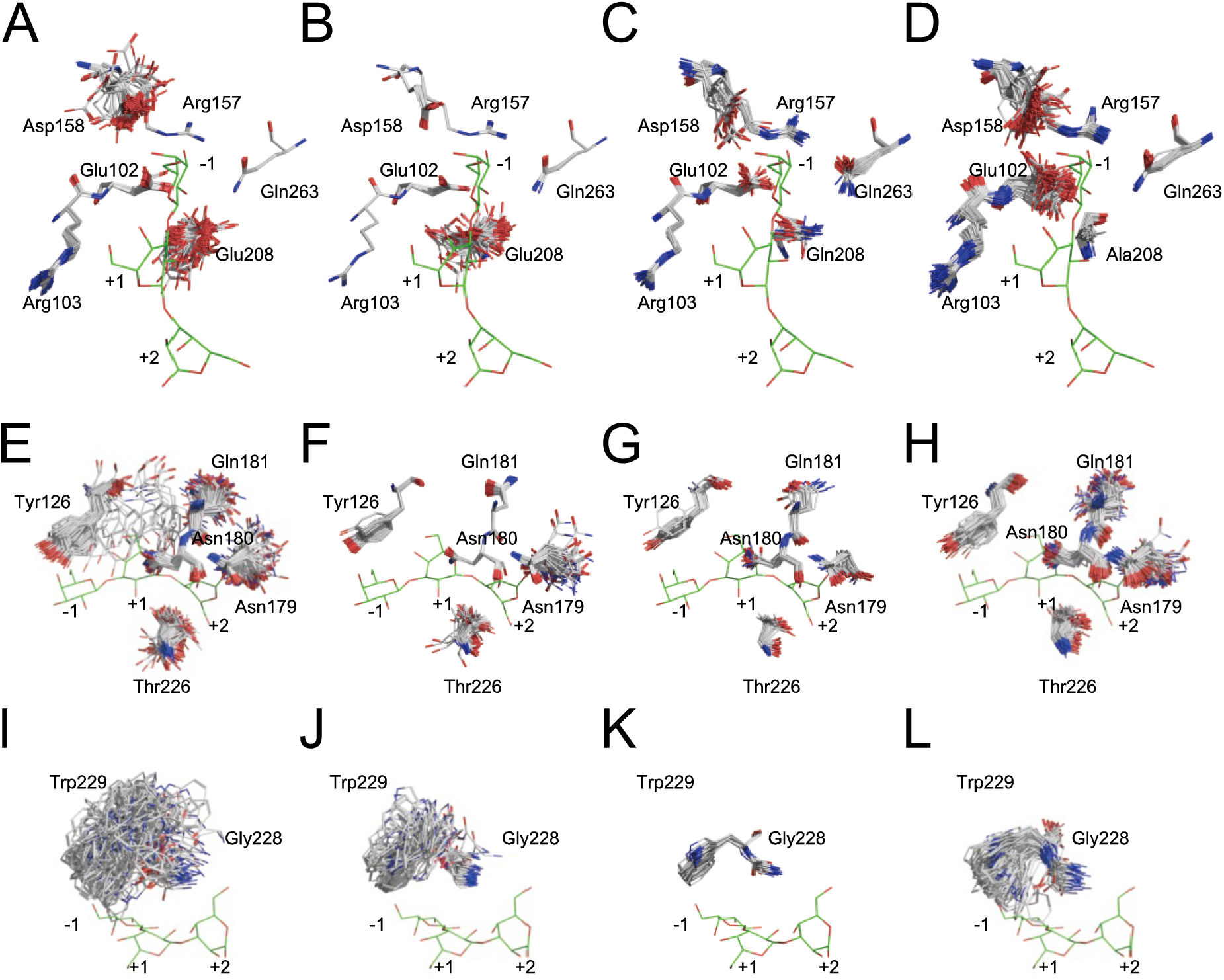
Results of ensemble refinement at the catalytic site. Each model is divided into three parts for clarity. A (E, I), B (F, J), C (G, K), and D (H, L) show WT, WT_Gal, E208Q_Gal3, and E208A_Gal3, respectively. Although WT and E208A_Gal3 contained multiple molecules in an asymmetric unit, the results obtained with multiple molecules were considered as an ensemble of one molecule in the present study. Letters indicate the chain names. Atoms are indicated in the same colors as Fig. 2. Gal3 of the structure of E208Q_Gal3 obtained by X-ray crystallography is arranged in each figure to maximize ease of comparison.

**Fig. 6.**
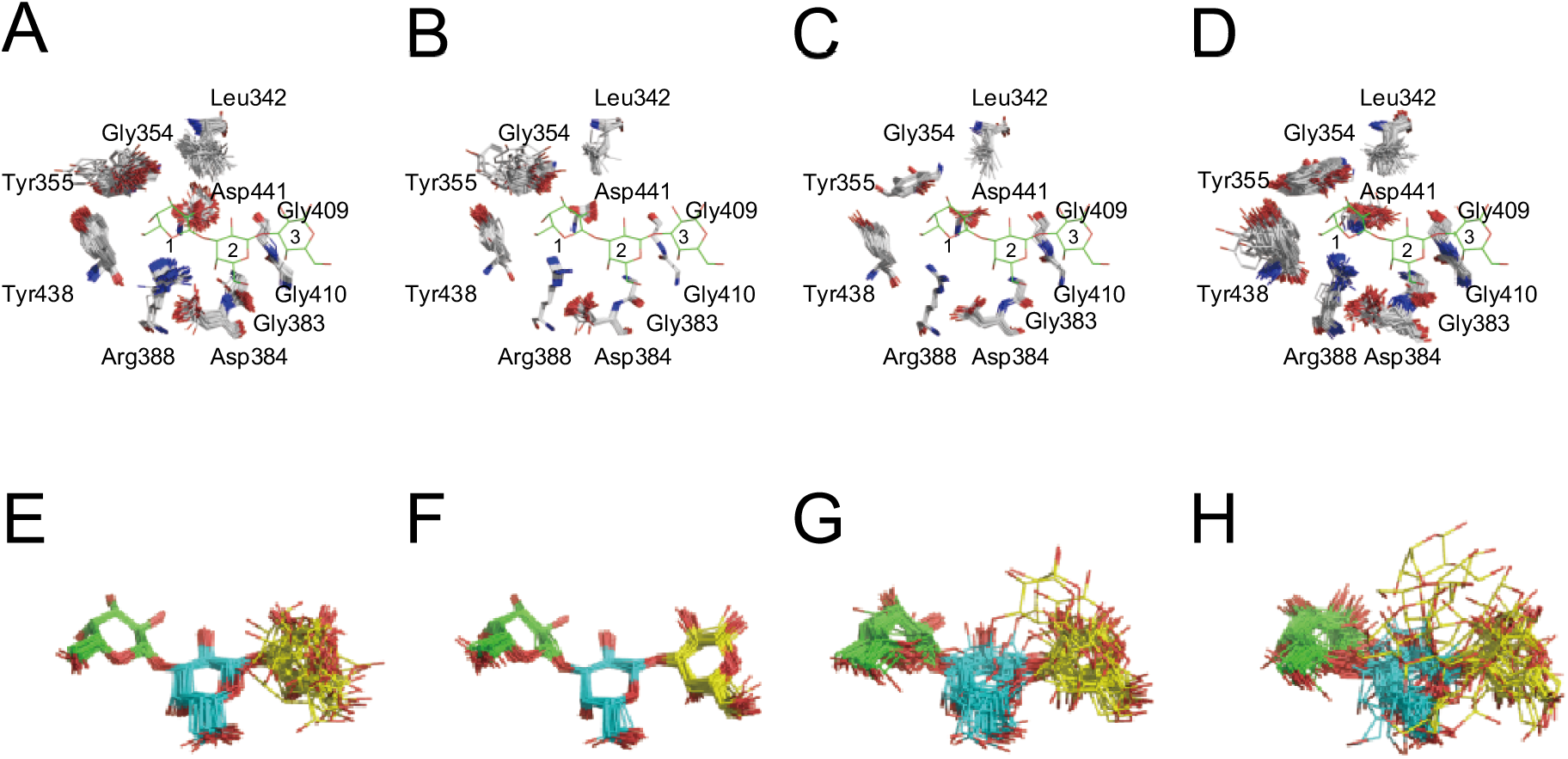
Results of ensemble refinement at the CBM ligand-binding site. A,B, C, and D: Residues related to ligand interaction. In this figure, Gal3 of chain A of refined E208A_Gal3 is drawn for comparison. E, F, G, and H: the ligands of each chain. Green, cyan, and yellow are used in order from the non-reducing terminal Gal. A and E, B and F, C and G, D and H represent chain A, B, C, and D of E208A_Gal3, respectively. Atoms are indicated in the same colors as Fig. 2.

**Table 3.**
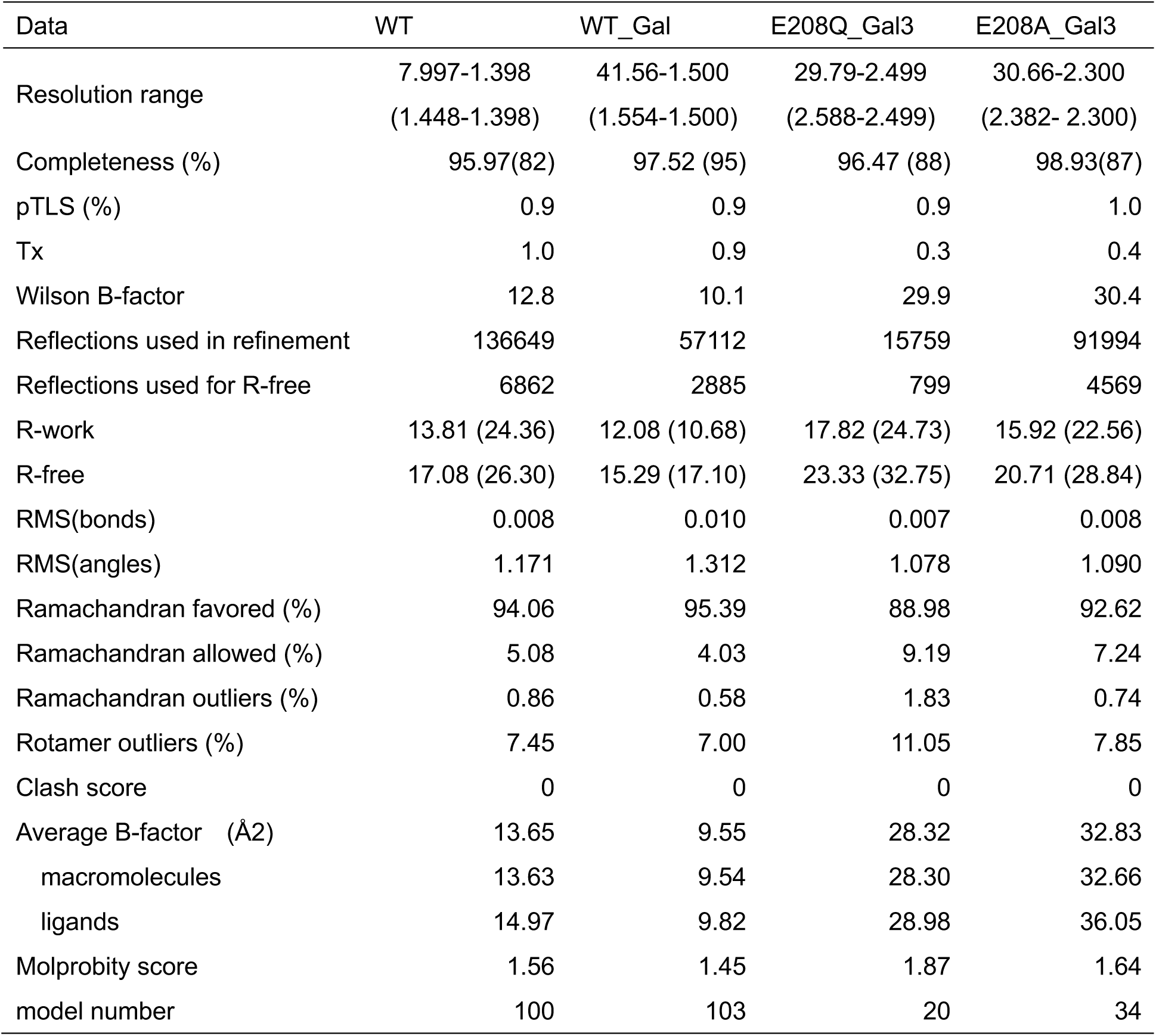
Refinement statistics of ensemble refinement

In the catalytic site, the vibration levels of some residues were significantly different between the apo and holo forms. As shown in Fig. 5, Tyr126, Arg157, Asp158, Asn179, Asn180, Gln181, Trp229, and Gln263 in the liganded structures (Fig. 5B, C, F, G, J, and K) showed smaller vibrations than in the apo structures (Fig. 5A, D, E, H, I, and L). These results indicate that side chain fluctuations converge upon ligand binding. Comparison of the Gal-bond structure (i. e. WT_Gal; Fig. 5B, F, J) with the Gal3-bond structure (i. e. E208Q_Gal3; Fig. 5 C, G, K) showed that the fluctuations of Glu(Gln)208, Asn179, and Thr226 of E208Q_Gal3 were smaller than these in WT_Gal. Therefore, it can be inferred that these residues recognize the ligands at the plus subsites. The catalytic acid, Glu208, has two major conformations in WT and WT_Gal. These two conformations were also reported in the BT3683 structure (12). Thus, the movement of this residue appears to be important for catalysis. Gln263 shows one conformation (Fig. 5A-D) that is identical to the result of the ensemble refinement of Asn92, known as imidic acid in *Pc*Cel45A (Fig. S3). Glu102 may distinguish non-reducing terminal Gal, since it interacts with the axial C-4 hydroxyl group of Gal_-1_ (12). The vibration degree of Glu102 was different between WTs and mutants, so its conformation does not depend on the ligand localization, but reflects interaction with Glu208, which serves as a general acid. Asp158 of WT and E208A_Gal3 show greater vibration than WT_Gal and E208Q_Gal3. The role of Asp158 is thought to be a pKa modulator, therefore its function and conformational stability might be related. Focusing on Fig. 5I-L, there were large differences in the fluctuation level of Trp229. E208Q_Gal3 (Fig. 5K) showed small movements of Trp229, but other structures showed much larger fluctuations (Fig. 5I, J, L). These results suggest that this Trp is normally flipped and forms a π-π interaction to anchor the ligand in the proper position upon arrival. A histogram of the dihedral angle is shown in Fig. S4.

As regards the ligand binding site of the CBM, a comparison of each chain of the E208A_Gal3 asymmetric unit showed no significant difference in the vibration levels of each residue involved in ligand binding (Fig. 6). However, ensemble refinement revealed that Gal_site 1 and Gal_site 2 do not show huge fluctuations, while Gal_site 3 has many conformations. They include the same conformation of each chain Gal of X-ray crystallography. Interestingly, a spatial difference in fluctuations was observed between ligands bound to the catalytic site and to the ligand binding site of CBM35 (Fig. 7). At the catalytic site, Gal_-1_ is anchored in the appropriate position, and Gal_+2_ appears to fluctuate in a planar fashion as it interacts with the surrounding residues. In the CBM, it was inferred that Gal_site 1 is fixed and Gal_site 3 is adsorbed at the appropriate location at the binding site while fluctuating in three dimensions.

**Fig. 7.**
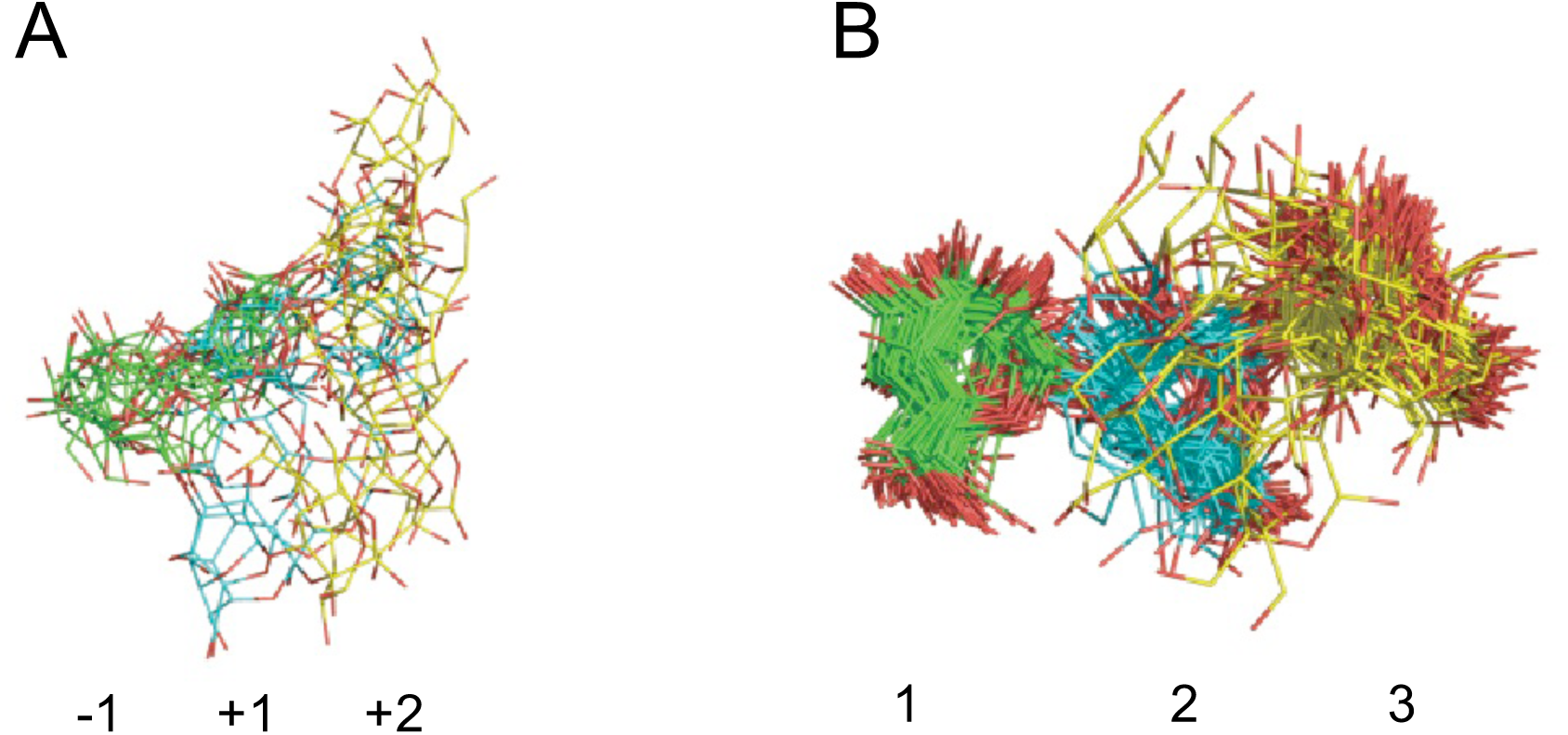
Ligand conformation of ensemble refinement at glance. A: ligand conformation of E208Q_Gal3 ensemble model. B: ligand conformation of E208A_Gal3 ensemble models with four chains aligned. Green, cyan, and yellow are used in order from the non-reducing terminal Gal.

## Discussion

Most exo-β-1,3-galactanases belonging to GH43_sub24 possess CBMs that can be classified into CBM35 or CBM13 (8). In this study, we elucidated the structure of a β-1,3-galactan binding module for the first time by solving the structure of a GH43_sub24 containing CBM35, and obtained the ligand-bound structures of both the catalytic and sugar binding domains of *Pc*1,3Gal43A. This is also the first study to reveal the structure of a eukaryotic exo-β-1,3-galactanase. This information will be useful to understand how the CBM35 module interacts with β-1,3-galactan in combination with the GH43_sub24 catalytic module.

### How does *Pc*1,3Gal43A hydrolyze β-1,3-galactan?

Although catalytic residues such as Glu and Asp are conserved in GH43 as a catalytic acid and base, respectively, GH43_sub24 lacks such a base residue. Mewis and co-workers suggested that GH43_sub24 may use Gln in the base role via conversion to imidic acid, or use an exogenous base, or utilize the Grotthuss mechanism of catalysis (8, 12). In this study, we measured the enzyme activity of six variants of the three residues speculated to be involved in the catalytic reaction. As shown in Fig. S1, production of Gal by the mutants was not detected by means of high-performance liquid chromatography (HPLC) analysis, suggesting that all three residues are essential for the catalytic activity of *Pc*1,3Gal43A. Glu102, Glu208, and Gln263 are speculated to serve in C-4 hydroxyl group recognition, as a catalytic acid, and as a catalytic base, respectively. These residues are well conserved in GH43_sub24, as shown in Fig. S5.

In GH43_sub24, only bacterial enzyme structures have been solved so far (http://www.cazy.org/GH43_24.html). In order to understand the catalytic mechanism of *Pc*1,3Gal43A, we compared its structure with those of BT3683 and *Ct*1,3Gal43A (Fig. 8). Most of the residues that interact with ligands are conserved in these three enzymes. In subsite -1, all residues, Glu57, Glu102, Arg103, Tyr126, Asp158, Glu208, Trp229, and Gln263, of *Pc*1,3Gal43A are conserved, indicating that the binding mode at subsite -1 is fully conserved in GH43_sub24. Based on the results of ensemble refinement, Trp229 showed huge fluctuation, especially in the apo structure (Fig. 5I-L). Trp541 of BT3683, which corresponds to Trp229 of *Pc*1,3Gal43A, has a polar interaction with Gal (12). Trp229 fluctuates in solution and plays a role in holding the substrate at the catalytic site through polar interactions. On the other hand, Asn179 and Thr226 of *Pc*1,3Gal43A are replaced by Asp490 and Cys538 in BT3683 and by Glu199 and Cys247 in *Ct*1,3Gal43A. Since all of these enzymes can accommodate a β-1,6-branched side chain (6, 12, 28), we considered that these residues are not related the mechanism of side-chain accommodation.

**Fig. 8.**
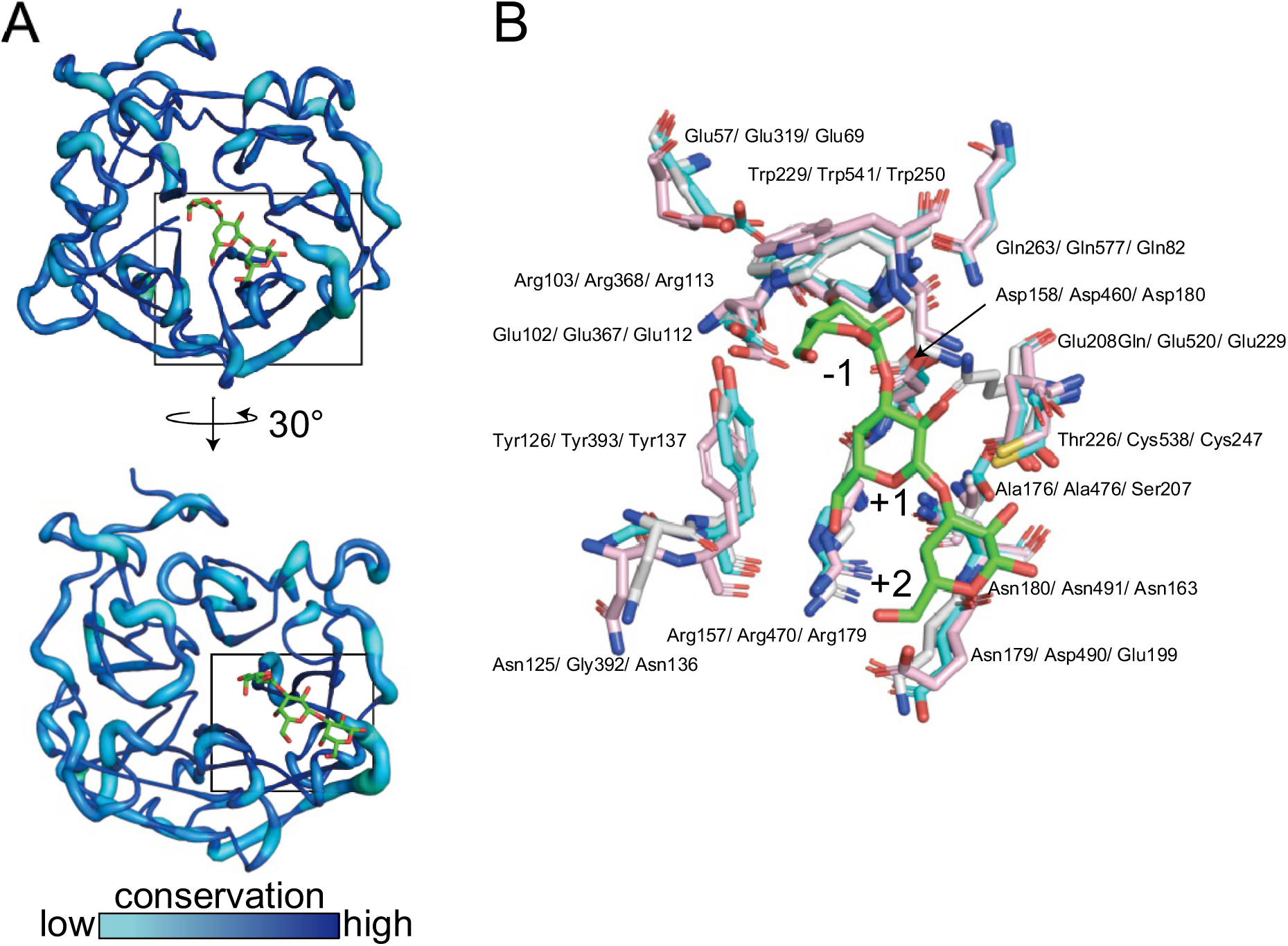
Catalytic domain structure comparison. A: Visualization of the degree of preservation of GH43_sub24. The degree of conservation of amino acid residues in the catalytic domain of GH43_sub24 was visualized using the ConSurf server (https://consurf.tau.ac.il), the query for BLAST was set to *Pc*1,3Gal43A, and the conservation degree was analyzed based on 150 amino acid sequences in the ConSurf server (47–51). The conservation degree is shown in graded color. Preservation degrees are shown in a gradient with cyan for the lowest degree of preservation and blue for the highest. B: Catalytic domain comparison of *Pc*1,3Gal43A and two GH43_sub24 galactanases. Catalytic center of E208Q_Gal3 of *Pc*1,3Gal43A (white, PDB ID: 7BYV), BT3683 (cyan, PDB ID: 6EUI), and *Ct*1,3Gal43A (pink, PDB ID: 3VSF). Red, blue, and yellow represent oxygen atoms, nitrogen atoms, and sulfur atoms, respectively. Residue names mean *Pc*1,3Gal43A/ BT3683/ *Ct*1,3Gal43A.

The bypass mechanism of *Pc*1,3Gal43A, which enables accommodation of the β-1,6-galactan side-chain so that the β-1,3-galactan main chain can be cleaved, appears to depend on the orientation of the C-6 methylol group of Gal3 at each subsite. The C-6 methylol group of Gal_-1_ is exposed to the solvent, so that the side chain can be accommodated externally. The C-6 methylol groups of Gal_+1_ and Gal_+2_ are also exposed to the solvent, so that the enzyme should be able to cleave the β-1,3-linkage of continuously β-1,6-substituted galactan, and a similar situation has been reported for BT3683 (12). Moreover, there are spaces near the non-reducing terminal Gal in these enzymes (12, 29). This enables the enzymes to degrade the main chain, even if the side chain contains multiple carbohydrates. Similarly, β-1,3-glucanases belonging to GH55 also bypass the β-1,6-glucan side chain and degrade β-1,3-glucan from the non-reducing end (29, 30). Comparing the surface structure of the catalytic site of *Pc*1,3Gal43A with that of these GH55 exo-β-1,3-glucanase from *P. chrysosporium* (*Pc*Lam55A), we see that *Pc*1,3Gal43A has a small pocket-like space capable of accepting the C-6 side chain of Gal at subsite -1 (Fig. 9A and B). In addition, the C-6 methylol group of Gals, located at the positive subsites of *Pc*1,3Gal43A, are exposed to solvent in a similar manner to that reported for SacteLam55A, GH55 exo-β-1,3-glucanase from *Streptomyce*s sp. SirexAA-E (Fig. 9A and C). Structures capable of accepting non-reducing terminal Gal with β-1,6-linked Gal are conserved among GH43_sub24 of known structure (Fig. 8 and S5). In the non-bypassing GH3 *Hypocrea jecorina* β-glucosidase (*Hj*Cel3A), the C-6 hydroxyl group of non-reducing glucose is oriented toward the enzyme, introducing steric hindrance (Fig. 9D; 31). In other words, enzymes bypassing side chains have a space adjacent to C-6 of the non-reducing terminal sugar, and the positive subsites are particularly wide, allowing side chains of the substrate to be accommodated. In contrast, enzymes unable to bypass the side chain have no space next to the -1 subsite and have a narrow entrance to the catalytic site, so that they are unable to accommodate side chains (Fig. 9D).

**Fig. 9.**
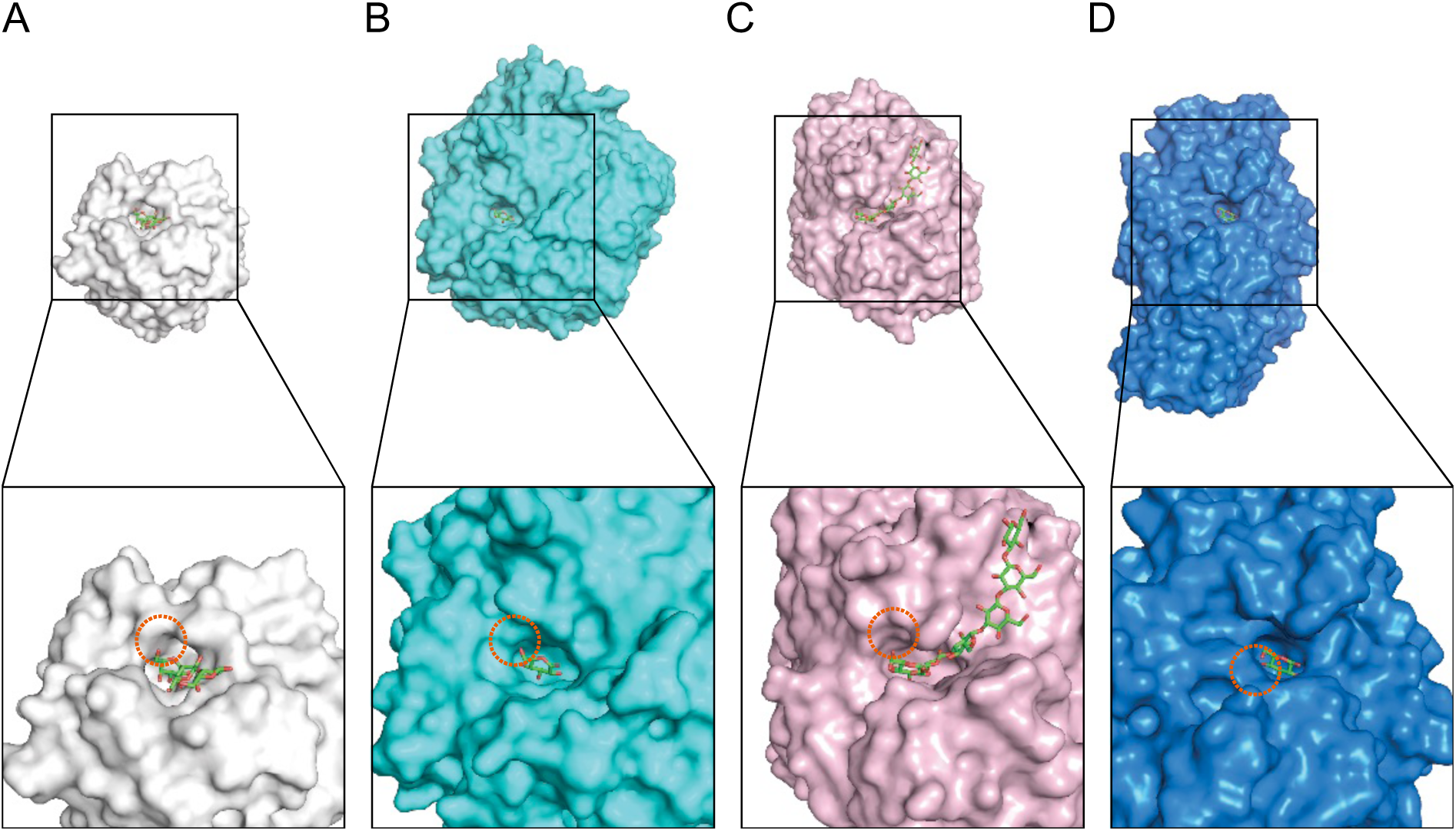
Structure comparison of the catalytic sites of *Pc*1,3Gal43A (A), GH55 exo-β-1,3-glucanase from *P. chrysosporium* (B; *Pc*Lam55A; PDB ID: 3EQO), GH55 exo-β-1,3-glucanase from *Streptomyce*s sp. SirexAA-E (C; SacteLam55A; PDB ID 4PF0), GH3 β-glucosidase from *H. jecorina* (C, PDB ID: 3ZYZ). A, B, and C hydrolyze the main chain of β-1,3-galactan or β-1,3-glucan, bypassing β-1,6-branched side chains (6, 29, 30). D hydrolyzes four types of β-bonds, and it does not bypass side chains (31, 52). The upper panel shows the overall surface structure and the lower panel shows an enlarged view of the catalytic region. Orange dashed circles indicate the space near the C-6 position of Gal or glucose at the non-reducing end.

Although the electron density of Gal3 was observed in the present study, *Pc*1,3Gal43A is proposed to have four subsites ranging from -1 to +3, based on biochemical experiments (6). As mentioned above, although *Pc*1,3Gal43A has a structure capable of accepting the C-6 side chain, its degradation activity towards β-1,3Λ,6-galactan is only approximately one-fifth that of the linear β-1,3-galactan (6). This difference in reactivity may be due to the structure of the sugar. The β-1,3-galactan in solution has a right-handed triple helical structure with 6 to 8 Gal residues per turn (32, 33), with the C-6 methylol group pointing outward to avoid collisions between the β-1,6-bonded Gal side chains (32). However, as shown in Fig. S6, Gal3 bound to the catalytic site of *Pc*1,3Gal43A is anchored to the enzyme, so that the helix of the glycans differs from the usual state in solution. Therefore, the reason why the hydrolytic activity of *Pc*1,3Gal43A towards β-1,3Λ,6-galactan is lower than that towards β-1,3-galactan may be interference between the β-1,6-Gal(s) side chains as a result of changes in the helical state of the main chain.

### How does *Pc*CBM35 recognize β-1,3-galactan?

Although the amino acid sequence similarity of CBM35s is not so high, important residues involved in ligand binding are well conserved (17). The modules belonging to CBM35 can be divided into four clades according to the mode of ligand binding, and the diversity in ligand binding and in the calcium ion-coordinating residue account for the various ligand binding specificities (17, Fig. 10A). Moreover, the residues involved in ligand binding of *Pc*CBM35 differ from those of CBM35, which binds to α-Gal of galactomannan. This CBM is one part of a protein predicted to be the β-xylosidase of *C. thermocellum* cellulosomal protein (Cte_2137; Fig. 10), which belongs to the same cluster as *Pc*CBM35 (17). There are some differences between the residues interacting with α-Gal of Cte_2137 and those interacting with β-Gal of *Pc*CBM35. For instance, the regions of Ala352 to Tyr355 and Tyr438 to Asp441 of *Pc*CBM35 correspond to Val39 to Gly42 and Ser136 to Asn140 of Cte_2137, which are related to ligand specificity (Fig. 10). Especially, Asn140 of Cte_2137 is not conserved but replaced Asp441 in *Pc*CBM35 and is located at the bottom of the ligand binding site. Furthermore, Trp108 of Cte_2137, which is conserved in *Pc*CBM35, plays a key role in stacking the pyranose ring (17), while in CBM35 of *Pc*1,3Gal43A, this Trp residue is replaced with Gly (Fig. 10B). In other words, although *Pc*CBM35 and Cte_2137 are in the same cluster, the residues involved in ligand recognition are different, and this difference affects the discrimination between β-Gal and α-Gal, and between galactan and galactomannan. It is still unclear how CBM35s acquire such variation of binding specificity within a similar binding architecture. However, a detailed understanding of the molecular mechanisms of polysaccharide recognition by CBM35 will be essential for efficient utilization of various types of biomass.

**Figure. 10.**
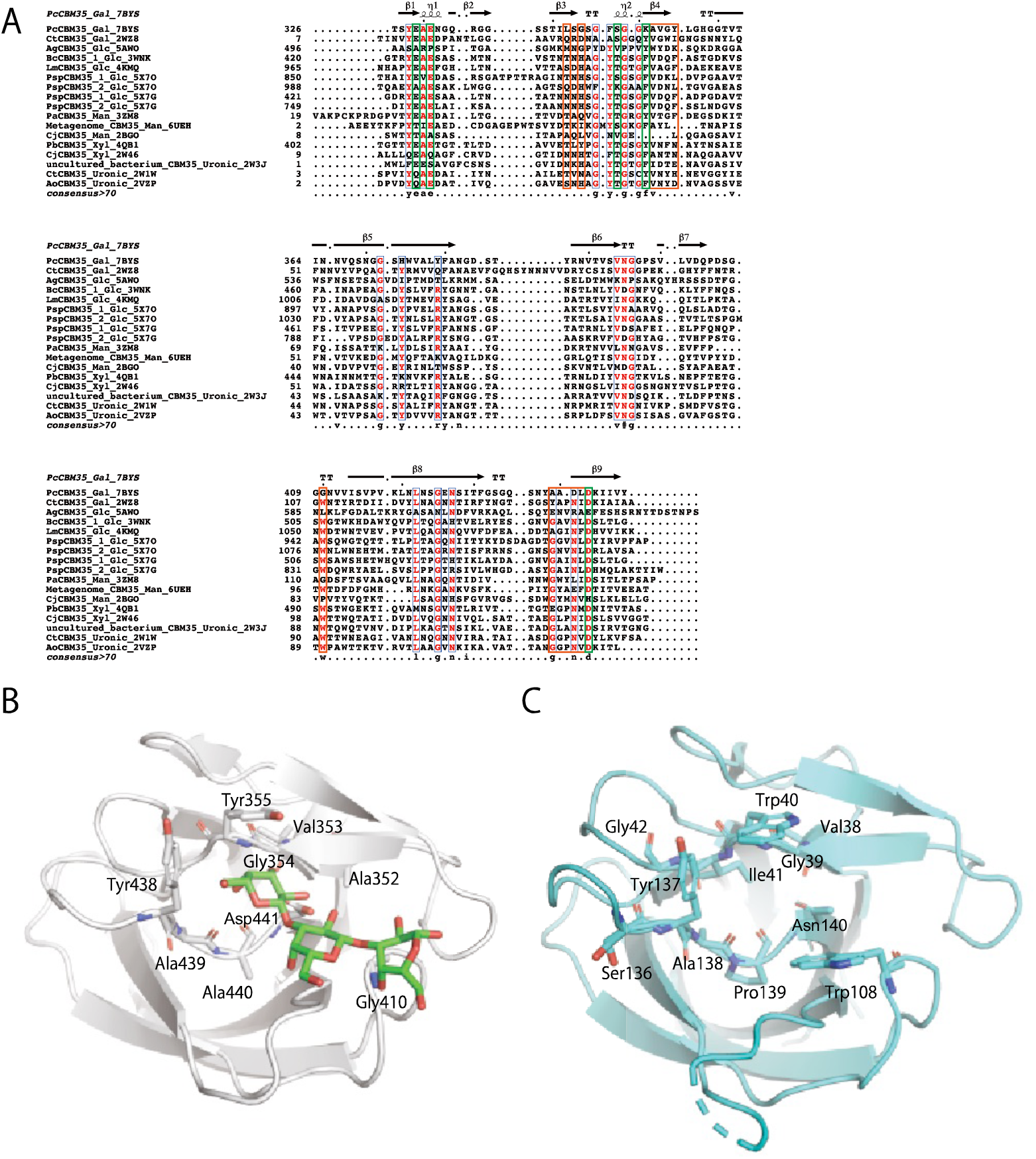
Sequence alignment of known CBM35s (A) and structure comparison between CBM35s of *Pc*1,3Gal43A (B) and Cte_2137 (C). A: Taxon names are shown as scientific names, ligand specificity and PDB ID only for brevity. When the same enzyme contains two CBM35 domains, the taxon name is indicated with 1 on the N-terminal and 2 on the C-terminal. Gal, Glc, Man, Xyl, and Uronic means ligand specificities for Gal, glucose, mannose, xylose, and glucronic acid and/or galacturonic acid, respectively. Among these, 3ZM8, 6UEH, and 2BGO, which bind to Man, are Type B CBMs, which show endo-type binding, while the other 14 are all Type C CBMs, which show exo-type binding. The alignment was built by using MUSCLE on MEGAX: Molecular Evolutionary Genetics Analysis (53, 54), and the figure was generated with ESPrint 3.0 (http://espript.ibcp.fr; 56). Orange and green boxes represent ligand binding and calcium ion binding residues, respectively. B and C: Ligand binding residues of *Pc*1,3Gal43A (chain A of E208A_Gal3) and Cte_2137 (PDB ID: 2WZ8). Red and blue mean oxygen and nitrogen, respectively. The green stick model represents Gal3.

In conclusion, we have determined the crystal structure of the catalytic and binding domains of *Pc*1,3Gal43A with the aim of reaching a detailed understanding of the mechanism of substrate accommodation by side-chain-bypassing galactanase. *Pc*1,3Gal43A uses Glu as the catalytic acid and Gln as the catalytic base, and has a structure in which the side chain of the substrate does not interfere with the catalytic reaction, thus making it possible to degrade the β-1,3-galactan main chain of AGPs despite the presence of the β-1,6-galactan side chain. Thus, although polysaccharides have a variety of molecular decorations, it appears that the structures of the degrading enzymes enable them to recognize specific features of the substrate while accommodating the variations. The introduction of mutations in substrate recognition residues to create enzymes with altered substrate recognition properties is expected to be helpful in the structural analysis of AGP glycans and also for the preparation of useful oligosaccharides.

## Experimental procedures

### Expression of *Pc*1,3Gal43A and its mutants

The E208Q, E208A, E102Q, E102A, Q263E, and Q263A mutants were constructed by inverse PCR using PrimeSTAR MAX (Takara, Tokyo, Japan). For crystallization, *Pc*1,3Gal43A WT, E208Q, and E208A from *P. chrysosporium* were expressed in *P. pastoris* and purified as previously reported (7). For reactivity assay, WT and mutants were purified by using SkillPak TOYOPEARL Phenyl-650M (c.v. = 5 ml, Tosoh, Tokyo, Japan) equilibrated with 20 mM sodium acetate buffer, pH 4.0, containing 1 M ammonium sulfate, and the enzymes were eluted with 20 mM sodium acetate buffer, pH 4.0, containing 0.7 M ammonium sulfate. SeMet-labeled *Pc*1,3Gal43A was expressed as previously reported (7).

### Preparation for β-1,3-galactooligosaccharides and crystallization of *Pc*1,3Gal43A

Gal2 and Gal3 were prepared as previously reported (6). The protein solution was concentrated to A_280_ of 10 ∼ 15 and used for the crystallization setup. The WT plate crystal used for data collection was obtained from a reservoir of 2.1 M ammonium sulfate, 0.1 M citrate buffer, pH 5.5. Other WT crystals were obtained from solutions in 16 % (w/v) polyethylene glycol (PEG) 10000, 0.1 M ammonium sulfate, 0.1 M bis-tris, pH 5.5, and 5.0 % (v/v) glycerol. SeMet crystals were obtained from 16 % (w/v) PEG 10000, 95 mM ammonium sulfate, 95 mM bis-tris, pH 5.5, and 4.8 % (v/v) glycerol. Two types of crystals, thin plate crystals (space group *P*2_1_) and rod crystals (*P*2_1_2_1_2_1_) appeared under the same condition. Cocrystallization of the E208Q mutant with 10 mM Gal3 in 16 % (w/v) PEG 10000, 95 mM ammonium sulfate, 95 mM bis-tris, pH 5.5, and 4.8 % (v/v) glycerol afforded thin plate crystals. The E208A mutant was cocrystallized with 10 mM Gal3 in 0.2 M potassium nitrate, 15% (w/v) PEG 6000, 20 mM sodium citrate, pH 4.5, and 5 % glycerol to afford bipyramidal crystals.

### Data collection and structure determination

Diffraction experiments for *Pc1,3Gal43A* crystals were conducted at the beamlines of the Photon Factory (PF) or Photon Factory Advanced Ring (PF-AR), High Energy Accelerator Research Organization, Tsukuba, Japan (Table 1). Diffraction data were collected using CCD detectors (Area Detector Systems Corp., Poway, CA, USA). Crystals were cryocooled in a nitrogen gas stream to 95 K. For data collection of the WT enzyme complexed with Gal3, *Pc*1,3Gal43A crystals were soaked in a drop containing 1 % (w/v) Gal3 for 10 min before the diffraction experiment. The data were integrated and scaled using the programs DENZO and SCALEPACK in the HKL2000 program suite (34)

Crystal structure was determined by means of the multiwavelength anomalous dispersion method using a SeMet-labeled crystal (7). Initial phases were calculated using the SOLVE/RESOLVE program (35) from five selenium atom positions. The resultant coordinates were subjected to the auto-modeling ARP/wARP program (36) in the CCP4 program suite (37), and manual model building and molecular refinement were performed using Coot (version 0.8.9, University of Oxford, Oxfordshire, England; 46), REFMAC5 (version 7.0.063, Science & Technology Facilities Council, England; 47), phenix.refine (40), and phenix.ensemble_refinement (27, 41, 42) in the Phenix suite of programs (version 1.13-2998-000, Lawrence Berkeley National Laboratory, USA; 51). The refinement statistics are summarized in Table 2.

For the analyses of WT, and ligand bound structures, structural determination was conducted by the molecular replacement method with the MolRep program (44) in the CCP4 program suite using the SeMet or ligand-free structure as the starting model. Bound sugars, water molecules and crystallization agents were modelled into the observed electron density difference maps. Calcium ion was modelled based on the electron density map and the coordination distances. Three N-glycans were observed, and the identified sugars were modelled. The stereochemistry of the models was analyzed with LigPlot + (version 1.4.5; 53, 54) and structural drawings were prepared using PyMOL (version 2.2.3, Schrödinger, LLC). The atomic coordinates and structure factors (codes 7BYS, 7BYT, 7BYV, and 7BYX) have been deposited in the Protein Data Bank (http://wwpdb.org/).

### Enzymatic activity assay of *Pc*1,3Gal43A and its mutants

To evaluate the reactivity towards Gal2 and Gal3 of WT and each mutant, 20 nM enzyme was incubated with 0.263 or 0.266 mM galactooligosaccharides in 20 mM sodium acetate, pH 5.0, for 30 min at 30 °C, respectively. The reaction was stopped by heating at 95 *°C* for 5 min. The supernatant was separated with 75% (v/v) acetonitrile on a Shodex Asahipak NH2P-50 4E column (Showa Denko, Tokyo, Japan), and the amount of released Gal was determined by HPLC (LC-2000 series; Jasco, Tokyo, Japan) with a Corona charged aerosol detector (ESA Biosciences, now Thermo Fisher Scientific Corporation, Massachusetts, USA). One unit of enzyme activity was defined as the amount of enzyme that releases 1 μmol of Gal per one minute per one nmol of enzyme under our experimental conditions.

## Acknowledgments

We would like to thank Dr. Takuya Ishida (Japan Aerospace Exploration Agency) for helping with the crystallization and structure refinement. We also thank the staff of Photon Factory for X-ray data collection.

## Author contributions

K. M., N. K., and Z.F. solved and refined structures; K.M. and N. S. assayed enzymatic activity; T. K. and Y. T. prepared galactooligosaccharides (Gal2 and Gal3); K. M., M. S, K. I, and S.K. wrote the manuscript, and all authors commented on the manuscript.

## Funding and additional information

This research was partially supported by a Grant-in-Aid for Scientific Research (B) 19H03013 to K.I.) from the Japan Society for the Promotion of Science (JSPS) and a Grant-in-Aid for Innovative Areas from the Japanese Ministry of Education, Culture, Sports, and Technology (MEXT) (No. 18H05494 to K.I.). In addition, K.I. thanks Business Finland (BF, formerly the Finnish Funding Agency for Innovation (TEKES)) for support via the Finland Distinguished Professor (FiDiPro) Program “Advanced approaches for enzymatic biomass utilization and modification (BioAD)”.

